# Assessment of population exposure to pesticide applications with a high-resolution map of pesticide use intensity for mainland France

**DOI:** 10.1101/2024.11.25.624818

**Authors:** Benjamin Nowak, Gaëlle Marliac

**Affiliations:** Université Clermont Auvergne, AgroParisTech, INRAE, VetAgro Sup, UMR Territoires, F-63370 Lempdes, France; Université Clermont Auvergne, INRAE, UMR GDEC, F-63000 Clermont-Ferrand, France

**Keywords:** Plant, Pesticides, Spatial analysis

## Abstract

While there is a lively debate about the potential harmful effects of pesticide use, the multiplication of data sources offers new ways of assessing these effects. To facilitate future studies, this article sets out a method for obtaining a map of pesticide use intensity for mainland France, taking into account the effect of each plot. To do so, a number of pesticide treatments was assigned to each plot based on statistical data concerning the average number of treatments per crop and per region. The final result is a raster with a resolution of 100m, which takes into account the average of treatments for the years 2019, 2020 and 2021. A first application conducted in this study showed that 25% of the French population was exposed to at least 1 pesticide treatment per year in their place of residence (5% exposed to at least 10 treatments per year). There were marked differences in exposure between regions, with particularly high exposure in the west and north of France, for example. The map of pesticide use intensity in France produced in this study is freely accessible for future studies.

## Introduction

The use of pesticides is a way to avoid crop losses due to biotic factors, but these products can also have deleterious effects on the environment and the human health. Along with other agricultural practices, the use of pesticides is one of the major causes of the decline in bird populations (Rigal et al., 2023), as well as on insect populations (Gandara et al., 2024). Recently, Frank (2024) showed that the decrease of insect-eating bat populations accross North America led to an increase in insecticide treatments, which in turn has led to an increase in infant mortality. Such studies depend on reliable and accurate quantification of pesticide use.

Today, the multiplicity of data available makes it possible to map agricultural practices on large scales, such as the adoption rate of cover crops (Nowak et al., 2022, 2021). Regarding crop protection, this type of approch may allow to assess the long-term effects of pesticides, beyond the traditional trials that precede the commercialization of a new active substance.

For France, at least two previous studies have sought to map pesticide use at country level. In 2022, the Solagro association published a map summarizing the average number of treatments per municipality, based on the surface area occupied by each crop and the average number of treatments per crop (Solagro, 2022). Yet there can be major differences within a single municipality, for instance between the residential areas bordering the fields and those in the city center.

Using a different approach, Guilpart *et al*. (2022) showed that 16% of French agricultural area was within 100m of plot boundaries. This buffer distance of 100m without pesticide applications is a compromise between the regulatory distance advocated by the French government (between 3 and 20m depending on crops, active substances and application methods) and the distance of 150m advocated by certain NGOs (Journal Officiel de La République Française, 2019; USDA, 2020). This previous study demonstrated that vineyards and orchards, two crops with a high number of treatments, were more frequently close to residential areas than field crops, but it lacked quantification of population exposure, in terms of the number of doses received or the number of people concerned.

This study aims to fill these gaps by (i) establishing a detailed map of pesticide applications across metropolitan France, and (ii) evaluating the percentage of the population residing in proximity to these treated areas, with a resolution of 100m. It provides a first quantification of exposure to pesticides in France. The maps produced are freely accessible, for possible use in future studies to assess the effect of pesticides on animal or human populations.

## Material and Methods

### Data processing description

Maps estimating the spatial distribution of pesticide applications have been produced for 2019, 2020 and 2021 (*Figure 1*). For this purpose, plot boundaries were defined using the French Land Parcel Identification System, which covers 99% of the French arable crop area (Agence de Services et de Paiement, 2024). In this file, a plot corresponds to an area cultivated with one main crop (or a crop mixture) in a given year.

**Figure 1.**
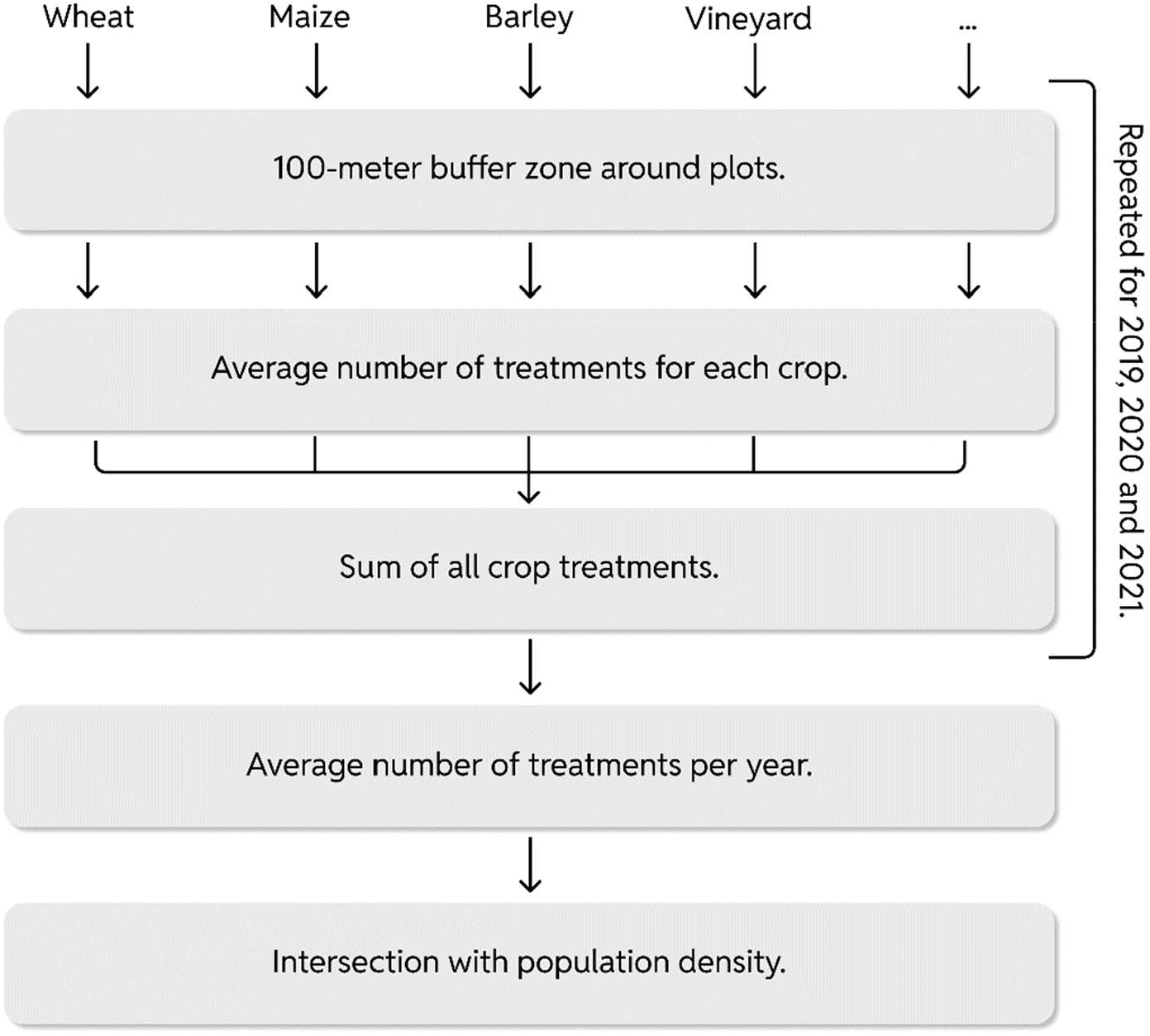
Summary of main data processing stages.

To take into account the diffusion of pesticides in the air, a positive buffer of 100m has been applied around plot boundaries. For each crop, the number of treatments applied has been defined on the basis of national crop practice surveys (Agreste, 2021, 2019, 2018). As the results of these surveys are aggregated on the scale of the former French administrative regions (22 regions for metropolitan France), the same number of treatments was assigned to all plots cultivated with the same crop in a given region.

Pesticide applications for all crops were then summed up to produce a map of pesticide applications for each of the years 2019, 2020 and 2021. In order to assess the number of people exposed to pesticide treatments, the resulting map of average annual pesticide applications was compared with the population density of the Global Human Settlement Layer project (Schiavina et al., 2023).

### Definition of the number of pesticide applications per crop

Surveys of cultivation practices carried out in France provide two values for each crop and region: the actual number of pesticide treatments applied and the Treatment Frequency Indicator (TFI). The latter is a standardized indicator that measures the quantity of pesticides applied to a plot in relation to the number of reference doses of the products used (Jorgensen, 1999). This second standardized indicator was used in this study.

### The case of organic farming plots

In order to differentiate between people exposed to synthetic pesticide treatments and those exposed only to pesticides authorized for use in organic farming, two different data treatments were carried out: the first taking into account all plots (whether conventionally or organically farmed), and the second taking into account only conventionally farmed plots. In the second case, organically farmed plots were removed before creating the pesticide application maps, using the database that records these plots (Agence Bio, 2024). This database include 80 to 85 % of the total number of plots under organic production, as not all plots under organic management are eligible for the European Common Agricultural Policy aid. When organic farming plots were included in the estimation of pesticide exposure, the average number of treatments affected per crop and per region was the same as for conventional farming, as the surveys have shown to be usually the case (Agreste, 2021, 2019, 2018). The main difference between the two production methods lies in the type of product used, rather than in the number of applications.

The data used in this study are summarized in *Table 1*. Data processing was realized through the Google Earth Engine platform (Gorelick et al., 2017). Analysis was conducted in R (R Development Core Team, 2009) and figures were produced using the package ggplot2 (Wickham, 2016).

**Table 1.** Datasets used for these studies.

| Source | Description | Data reference year |
| --- | --- | --- |
| Agence de Services et de Paiement (2024) | Boundaries of the plots (conventional or organic farming) declared under the European Common Agricultural Policy | 2019, 2020 and 2021 |
| Agence Bio (2024) | Boundaries of the plots under organic production declared under the European Common Agricultural Policy | 2019, 2020 and 2021 |
| Agreste (2021) | Treatment Frequency Indicator for field crops | 2021 |
| Agreste (2019) | Treatment Frequency Indicator for vineyards | 2019 |
| Agreste (2018) | Treatment Frequency Indicator for fruit production | 2018 |
| Schiavina et al. (2023) | Population density | 2020 |

## Results and discussion

### Map of pesticide applications

The map of pesticide use in mainland France was obtained by combining plot boundaries with the average number of treatments per crop (*Figure 2*).

**Figure 2.**
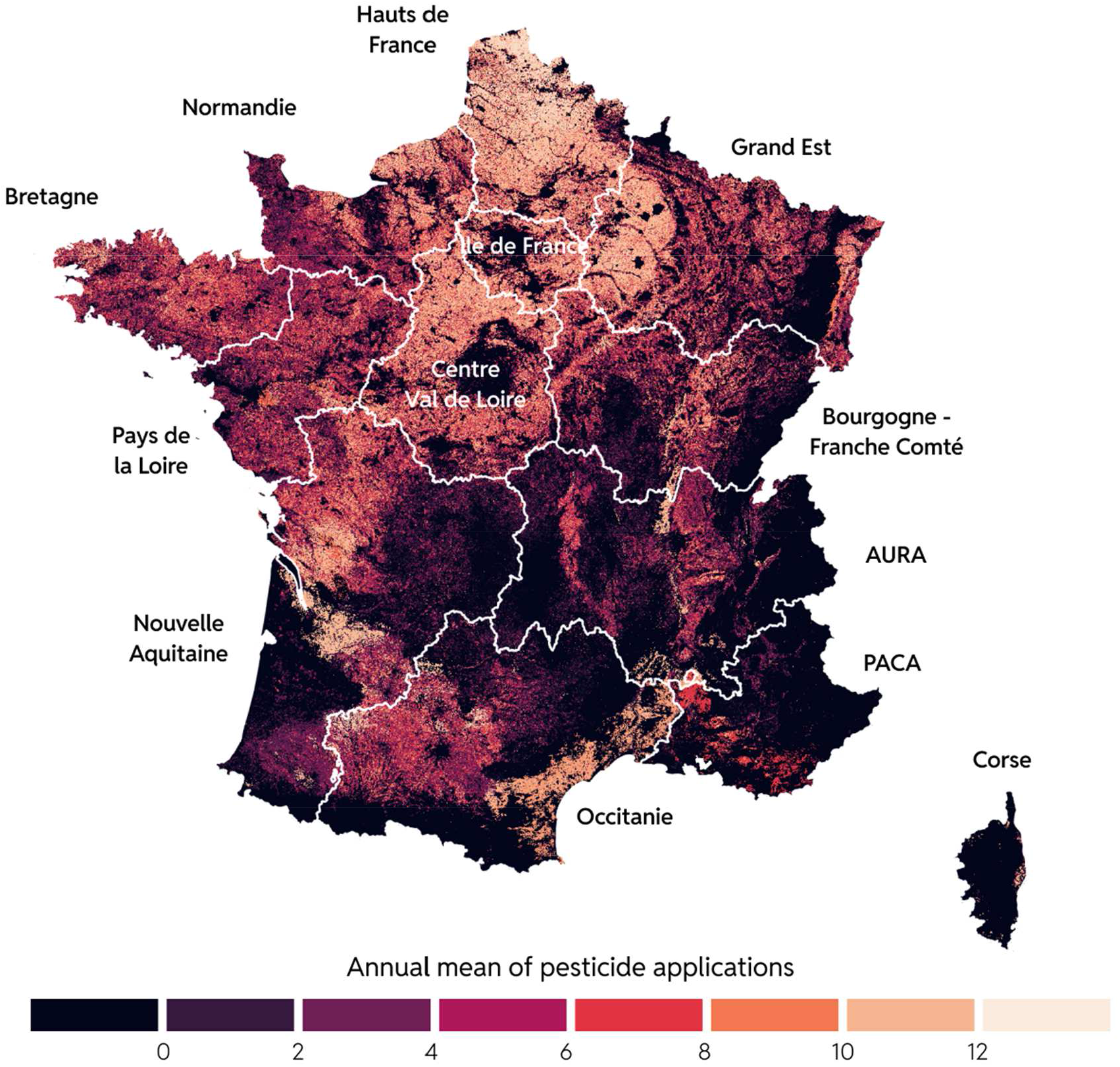
Map of estimated annual Treatment Frequency Indicator for the period 2019 to 2021. White lines represent the administrative boundaries of the new French regions.

This map shows the wide diversity of pesticide use across the region. The first factor of diversity concerns the distribution of agricultural land in the country (e.g. few agricultural plots in the main cities or natural areas). The second factor concerns the distribution of crops, with certain crops such as vines and arboriculture showing higher numbers of treatments than most field crops, for example. The third factor concerns the variability between regions in the number of treatments for the same crop.

The combination of these factors explains in particular the high Treatment Frequency Indicator in the Hauts de France region, which is a region with a large agricultural area, crops such as beet or potatoes with a high number of treatments (e.g. TFI between 24 and 25.4 for potatoes in northern France). but also, for a given crop, a higher number of treatments than regions located further south (for example, between 8.5 and 9.8 treatments for winter soft wheat in Hauts de France compared with between 3.8 and 4.7 for the same crop in the AURA region).

### Population exposed to pesticide applications

Population exposure to pesticide treatments was obtained by combining the map of treatments with the map of population density, as shown in *Figure 3*. Different patterns can be distinguished on the map, depending on the intensity of pesticide use and the organization of urban areas. These disparities will be discussed in the next section.

**Figure 3.**
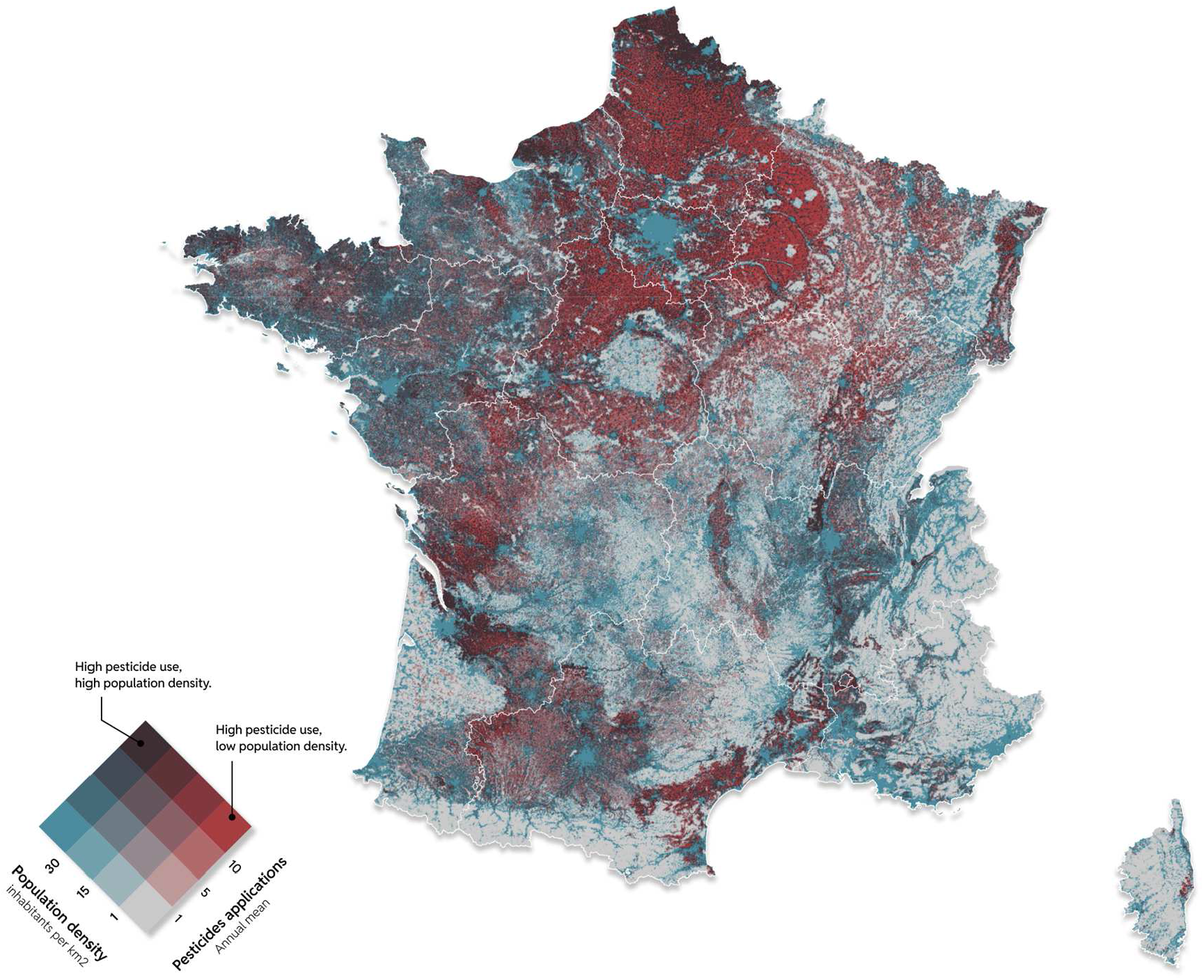
Bivariate choropleth of pesticide use intensity and population density.

Overall, approximately 24.6 % of the French population is exposed to at least one pesticide application per year less than 100 meters from their place of residence (*Figure 4*).

**Figure 4.**
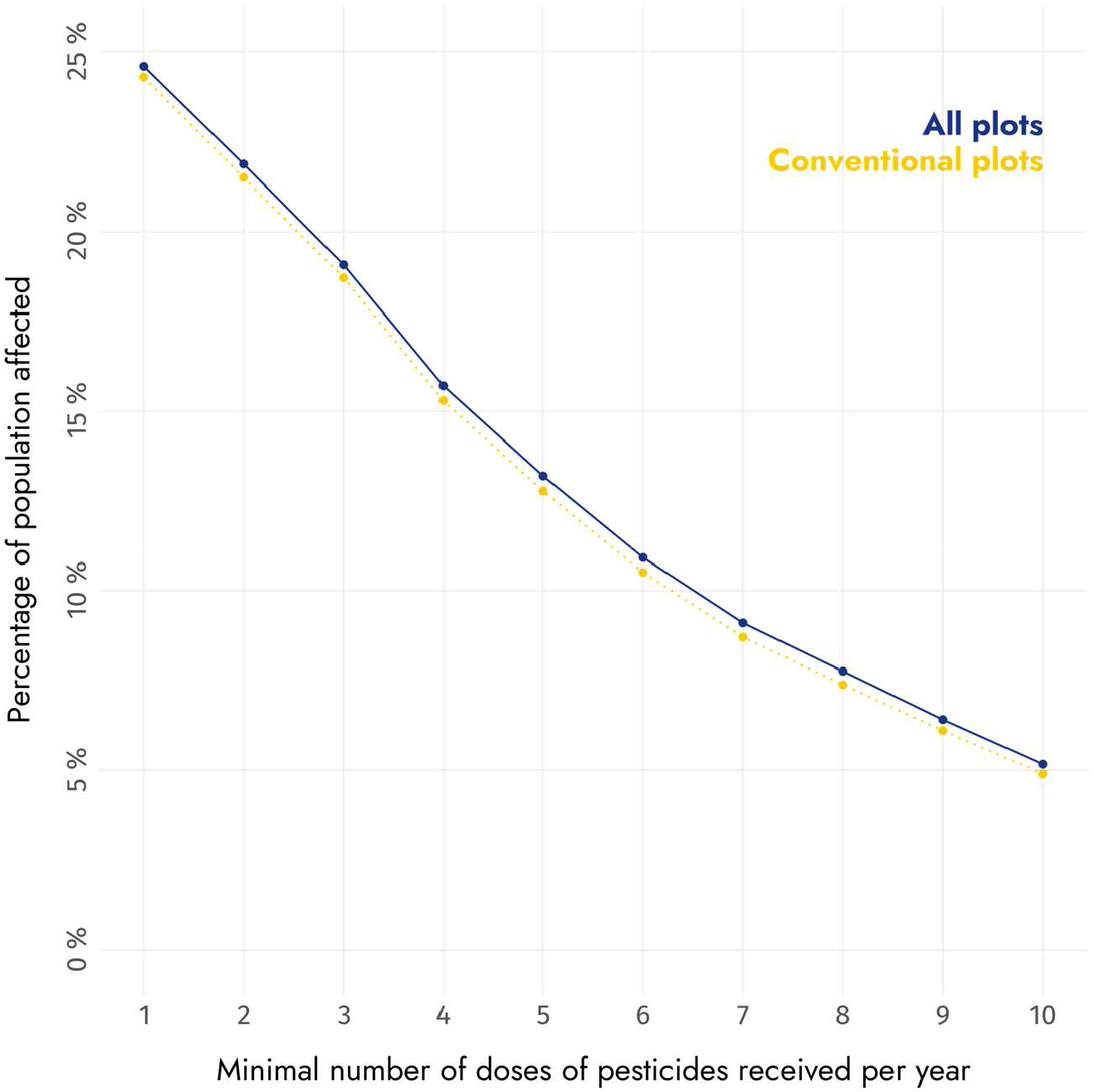
Percentage of the population of metropolitan France exposed to pesticide treatments for the period 2019 to 2021. The solid curve represents the estimate for all plots (conventional or organic), while the dotted curve represents exposure to conventional pesticides only.

This figure decreases only to 24.3 % when application of pesticide on organic farming plots are ignored. Such a result is partly explained by the relatively low proportion of organic farming in France, which represents only 10.4% of the agricultural area (Agence Bio, 2024). The ratio of organically farmed areas is even lower for field crops alone (5.2% in 2020).

But the small difference between total exposure and exposure to synthetic pesticides only may also be due to the spatial proximity of organic and conventional plots, which means that homes close to agricultural areas are often surrounded by plots cultivated with these two types of production.

Finally, a third explanatory factor may lie in the geographical distribution of conventional and organic plots, with organic plots more likely to be found in areas with lower yield potential, while conventional plots remain in more productive areas, which are generally close to housing (cereal plains, etc.).

The proportion of the population affected decreases when considering a larger number of doses, with for example 13.2 % and 5.2 % of the population exposed to 5 and 10 conventional or organic pesticides applications, respectively.

A further improvement of this map could be to distinguish the type of pesticides used according to their level of toxicity (Tao et al., 2020).

### Spatial variability of exposure to pesticides

As already suggested in *Figure 3*, residents’ exposure to pesticides varies widely from region to region (*Figure 5*).

**Figure 5.**
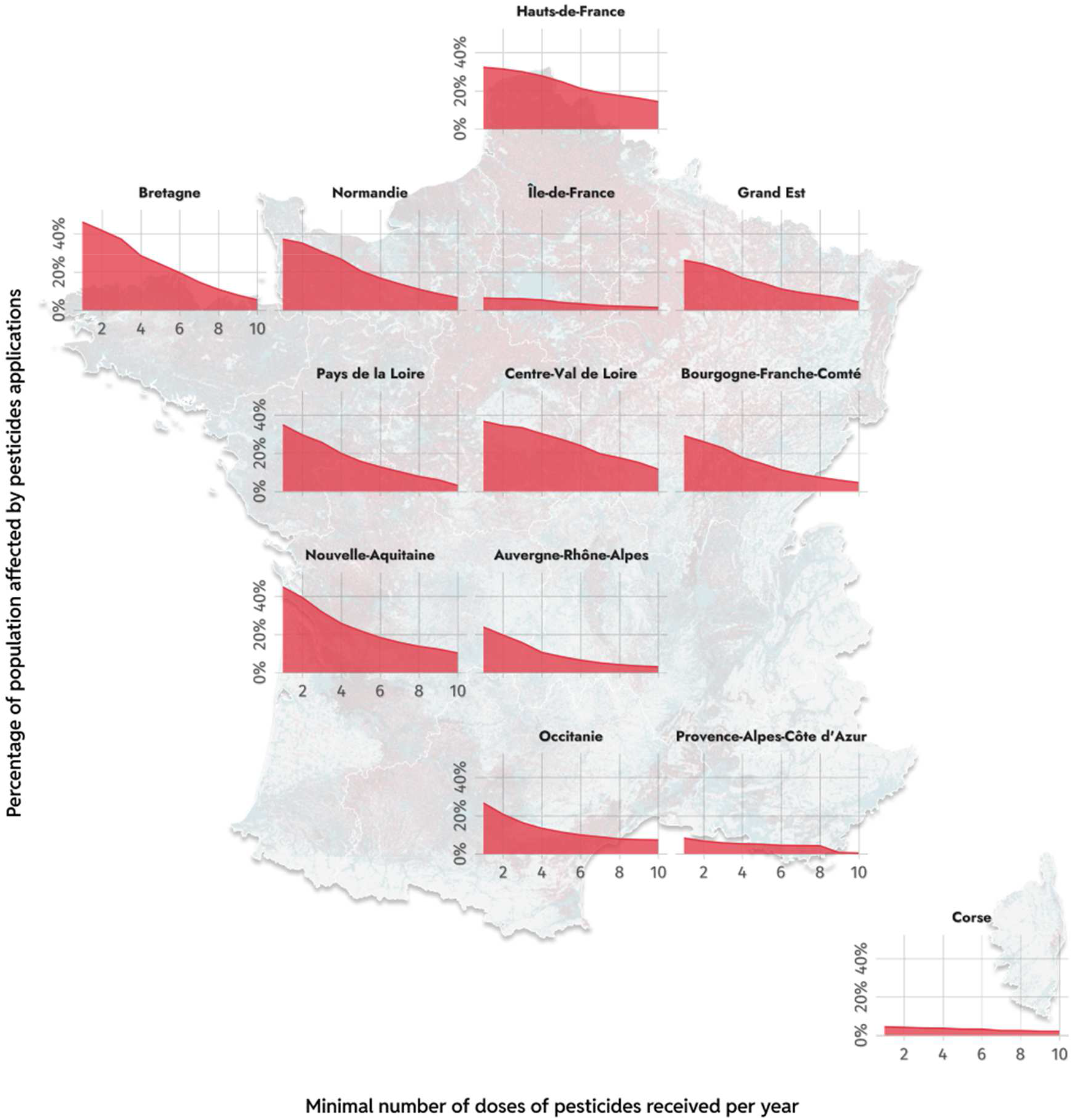
Spatial variability of pesticide exposure between regions. The x-axis shows the number of minimum doses applied (within 100m of the place of residence), while the y-axis shows the percentage of the region’s population affected.

In a highly urbanized region like Île de France, around Paris, exposure is very low. This is also the case in hilly regions with a high proportion of natural environments, such as Corsica.

Conversely, regions with a high level of agricultural activity, such as Brittany or Centre-Val de Loire, have higher levels of exposure, with around 40% of inhabitants exposed to at least one annual pesticide treatment within 100m of their place of residence. Such a result can be explained by the proximity of housing to fields in these regions, with many small urban areas intermingled with agricultural areas.

Some regions, such as New Aquitaine and Hauts de France, also feature a high proportion of inhabitants exposed to at least 10 treatments, which can be explained by the high number of crop treatments in these regions (e.g. vineyards in New Aquitaine).

## Conclusion

The map of pesticide use intensity produced in this study takes into account in a spatially explicit way the influence of each plot of the French Land Parcel Identification System. It is therefore more precise than the Adonis map, which was previously the reference in France, and for which pesticide use was aggregated at commune level (Figure 6).

**Figure 6.**
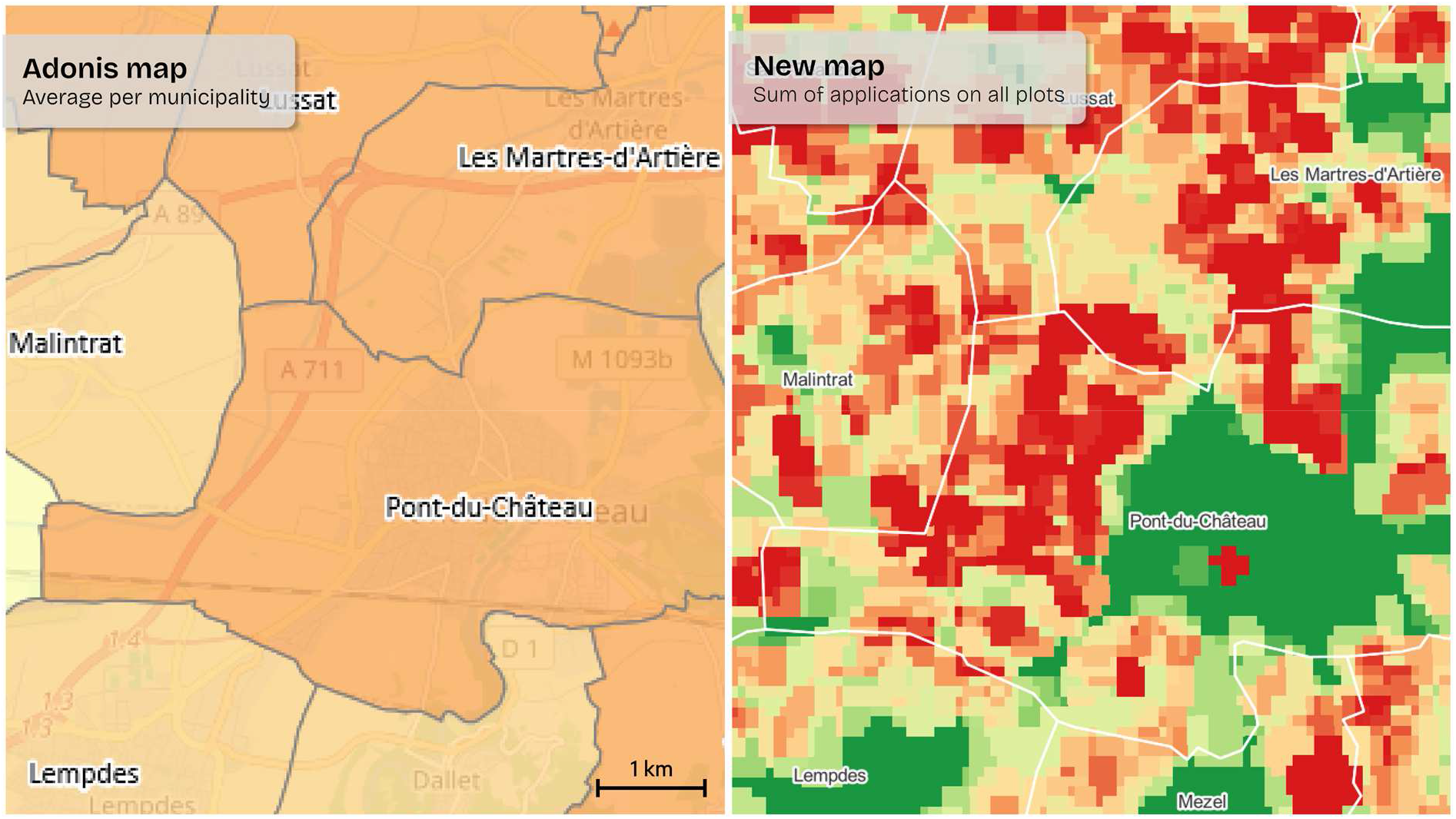
Comparison of Adonis map with the map produced in this study for a few municipalities of the Puy-de-Dôme (63) department. In both cases, the color scale represents the number of pesticide applications (increasing from green to red).

This increase in precision makes it possible, for example, to accurately assess the population’s exposure to pesticides, as presented in this study. The maps produced as part of this study are freely accessible to support other future applications, such as those related to the evalution of the impact of pesticides on human health or the evolution of wild animal populations.

## Data availability

The maps of pesticide treatments in mainland France created for this study and presented here are available for download at this link : https://zenodo.org/records/14202332

The average treatment map may also be consulted online, at this link : https://bjnnowak.quarto.pub/exposition_pesticide/

